# Ultrafast contour imaging for time-domain diffuse optical tomography

**DOI:** 10.1101/2020.09.06.285437

**Authors:** Xiaohua Feng, Liang Gao

## Abstract

Diffuse optical tomography (DOT) is well known to be ill-posed and suffers from a poor resolution. While time domain DOT can bolster the resolution by time-gating to extract weakly scattering photons, it is often confronted by an inferior signal to noise ratio and a low measurement density. This is particularly problematic for non-contact DOT imaging of non-planar objects, which faces an inherent tradeoff between the light collection efficiency and depth of field. We present here ultrafast contour imaging, a method that enables efficient light collection over curved surfaces with a dense spatiotemporal sampling of diffused light, allowing DOT imaging in the object’s native geometry with an improved resolution. We demonstrated our approach with both phantom and small animal imaging results. ©2020 Optical Society of America

---

Diffuse optical tomography [1–5] can probe non-invasively deep into scattering medium like biological tissues and provides quantitative measurement of the optical absorption and scattering properties. Though extensively employed both in the pre-clinical research and clinical imaging, it is plagued with a poor spatial resolution due to the ill-posed inversion of photon transport in the diffusive regime. Improving the spatial resolution, therefore, has been a central task of DOT over the past decades.

An effective approach to this end is to perform time-resolved measurements of the diffused light and extract the weakly scattered photons (i.e. early photons) through time-gating. Imaging with early photons markedly reduces the ill-posedness of the inversion problem, thereby enhancing the resolution [6–10]. Using time-gating, millimeter resolution had been demonstrated by time domain DOT (TD-DOT) on small animals [8,11]. Better resolution could be achieved in principle by using shorter time gates. However, the early photons also become exponentially weaker in this case, making it challenging to collect enough photons for a stable image reconstruction. The second approach is to sample the diffused photons more densely, as exemplified by the high-density DOT systems [12–16], which considerably improved the localization accuracy in human brain imaging [14,15]. Such improvement is possible because the spatial sampling in previous DOT systems were significantly below the Nyquist sampling rate for a targeted reconstruction resolution. To push the imaging resolution of DOT further, a natural solution is therefore to combine time gating with a high density sampling. This, however, faces a few technological challenges.

The first is the acquisition of a high-density TD-DOT data with a temporal resolution down to tens of picoseconds. Most previous TD-DOT systems relied on a single photon avalanche diodes (SPAD) or photo multiplier tube (PMT) [6] with a time-correlated single photon counter (TCSPC), which requires exhaustive temporal scanning to obtain a gray-scale waveform. A dense measurement will mandate either a large number of channels, which is cost prohibitive, or an extensive spatial scanning, which is time-consuming. While parallel ultrafast detectors like intensified CCD cameras can achieve dense spatial sampling, it still need time stepping and the temporal resolution is typically limited to 200 ps, yielding a sparse temporal data once time-gated.

The second challenge is to collect enough early photons for a stable reconstruction. In both fiber-based and non-contact DOT systems [17], the light collection is dictated by the sensor size and detection numerical aperture (NA) on the object side. While the fiber based approach can readily obtain a large NA, it is inherently difficult to obtain a dense measurement of time-resolved data. On the other hand, the detection NA of non-contact methods is often compromised for an increased depth-of-field to accommodate non-planar objects. Large defocus would introduce crosstalk between adjacent detection channels and ultimately degrade reconstruction accuracy and resolution. Previous efforts for minimizing defocus includes restricting the field of view [18], using a small detection numerical aperture [19], or fitting the object into a planar shape[8], which are either tedious to apply or sub-optimal for data collection. Though limiting the detection NA can increase the depth of field proportionally, the resultant quadratic reduction of collected light makes short time-gating impractical.

To address above challenges, we present here ultrafast contour imaging (UCI) for non-contact TD-DOT, which employs a spatial light modulator (SLM) for imaging over a curved surface and a streak camera for dense spatiotemporal sampling of diffused photons. The capability to focus onto a surface conforming to the object contour enables a large detection NA to be used for efficient light collection, thereby allowing short time-gating for resolution improvement.

The ultrafast contour imager, illustrated in Fig. 1(a), employs a streak camera to parallelly record the diffused photons emanating from the object. To image the object in its native geometry without defocus error, we captured the 3D shape of the object with a depth camera and then synthesize a 2D phase map *φ*(*x, y*) on a high resolution SLM to adjust the imaging plane of a collection lens onto a surface conforming to the object’s contour, as indicated in (b). This allows high density data collection over a large field of view. The phase map *ϕ*(*x, y*) effectively function as a diffractive lens with spatially-varying focal lengths *f*(*x, y*), which is calculated for each measured object contour via ray tracing: by casting the chief ray from each 3D point of the object to the streak camera, one can locate the unique intersection pixel on SLM, and with a known system geometry, the needed focal length for refocusing from the nominal object plane onto the 3D point can then be computed and assigned to that pixel.

The key for achieving a spatially-varying focal length is to note that the deflection of a light ray by a phase map is only determined by the local phase around the impinging point [20]. To make the patch around pixel (*x, y*) to function as a lens with a focal length of *f*(*x, y*), the phase function *ϕ*(*x, y*) need to satisfy:

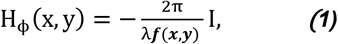

where *λ* is the working wavelength, ***I*** is an identity matrix and ***H*** denotes the Hessian operator. Such *ϕ*(*x, y*) exists exactly only when *f*(*x, y*) is a constant. For a spatially-varying *f*(*x, y*), only an approximate solution can be obtained by solving the following least square problem [20]:

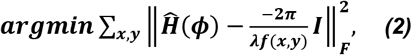

where 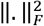 is the Frobenius norm. The least-square approach naturally promotes smooth solution for *f*(*x, y*), which is not an issue for two reasons: firstly, most objects lack sharp transitions in their 3D shapes (hence smooth) and secondly, the focal surface still possesses a finite depth of field to tolerate residual defocus error.

**Fig. 1.**
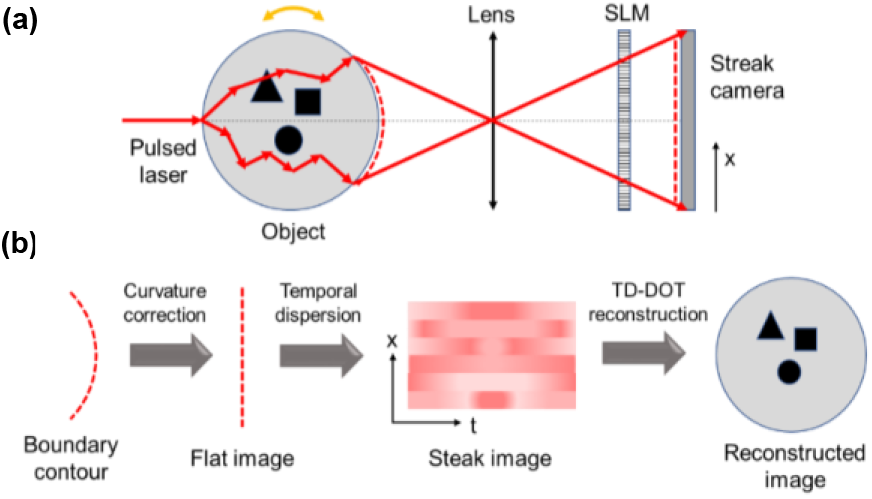
Operating principle of UCI-DOT. (a) A collimated laser pulse is transmitted through the object and the photons are collected over the boundary by a compounded lens (lens and SLM) and fed to the streak camera. (b) To focus on a curved boundary, the measured contour was transformed into a flat focal plane by synthesizing a phase map on SLM. Temporal DOT signals are then recorded as a streak image.

The system implementation is shown in Fig. 2(a). An 808 nm femtosecond laser (~2 uJ pulse energy and a pulse length of 200 fs) is collimated with a diameter of 1 mm and incident onto the object along the optical axis of the imaging lens (an achromatic doublet pair, Thorlabs, MAP105075-B, with 50 mm and 75 mm focal length). The object is placed about 50 mm away from the collecting lens, leading to a detection NA ~0.25 on the object side. A projector (LightCrafter3500, Texas Instruments) illuminates the objects obliquely and forms a structured-light stereo pair with a 2D CMOS camera (Blackfly S USB3, FLIR Inc.), which measures the 3D shape of the object with sub-millimeter accuracy. Afterwards, the phase map is synthesized and loaded into the reflective phase-only SLM (GAEA, Holoeye Inc.) to enable parallel measurements over the object boundary by the streak camera. The field of view of streak camera on the object side is ~12 mm and the pixels are binned to provide 60 effective detection points on the object. To acquire a dense time-resolved DOT data, the object is mounted on a 3D printed holder and rotated by a motorized stage over 360 degrees at an angular step of 6 degrees. The workflow of depth-mapping, phase synthesis and data acquisition, as summarized in Fig. 2(b), is repeated at each step before a final 2D image reconstruction.

**Fig. 2.**
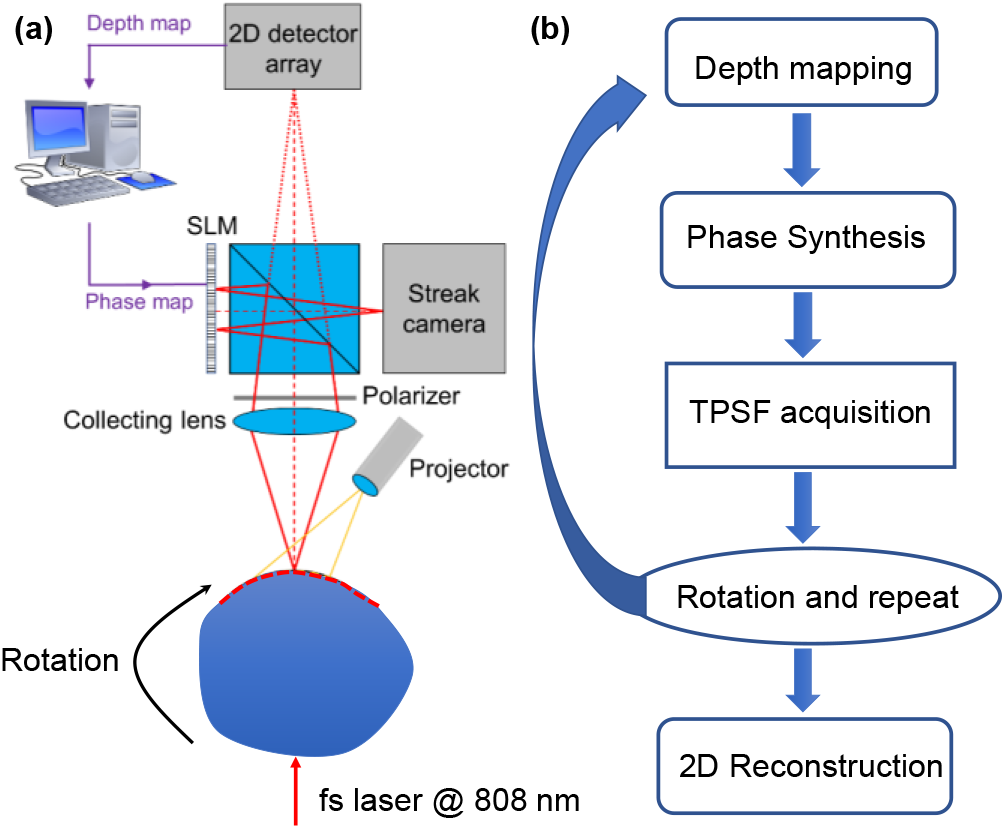
System schematic. (a) The structured-light depth camera (projector and camera) captures the contour of the object at each rotation angle and guides the synthesis of the phase map to be displayed on the SLM. The streak camera records parallelly 60 detections points (after binning) on the object along one dimension over a field of view of 12 mm. (b) Workflow of the system for 2D time-domain DOT imaging.

The resultant dataset contains, for a single 2D slice of the object, 60 (angular) × 60 (spatial) measurement pairs with a temporal resolution of 30 ps over a time window of 5 ns, yielding a spatiotemporal sampling of 3600 (spatial) ×150 (temporal), the densest dataset even when compared to those in 3D TD-DOT imaging systems. Though it is also feasible to scan the streak camera to sample the remaining spatial axis for 3D reconstruction, which offers best imaging retrieval owing to the inherent 3D light diffusion [21,22], acquiring such a huge dataset is time-consuming and the resultant reconstruction also becomes computationally challenging. As a result, we confine this study to 2D imaging.

For image reconstruction using the finite element method (FEM) [23], we first recover the 3D shape of the object by registering the object contour measured at all the angles and then generate a fine 2D mesh for the acquired slice. The data type used for TD-DOT reconstruction is the recorded temporal waveform, which is truncated when time gating is applied. As time-stepping in FEM is computationally intensive, we accelerate the image reconstruction using Gauss-Newton method [24] on a graphical processing unit (GPU).

Figure 3 demonstrate the proposed contour imaging over a slanted surface that shows large depth variations of ~8 mm. Without applying the contour imaging (by displaying a constant 0 phase map on SLM), only the central part of the image is in focus and thus resolving the letters (1 mm in height) in Fig. 3(a). In contrast, after applying contour imaging, the resolvable field of view in the resultant image in Fig. 3(b) is substantially enhanced (about 5 times), despite of some aberrations at the peripheral region. The dominant aberration is astigmatism because the diffractive lens on SLM shows a spatially-varying focal lengths *f*(*x, y*). Although astigmatism reduces effective image resolution, TD-DOT can tolerate a virtual sensor size comparable to its reconstruction resolution, which is on the order of 1 mm. As a result, the proposed contour imaging alleviates defocus error below astigmatism for efficient DOT signal acquisition.

**Fig. 3.**
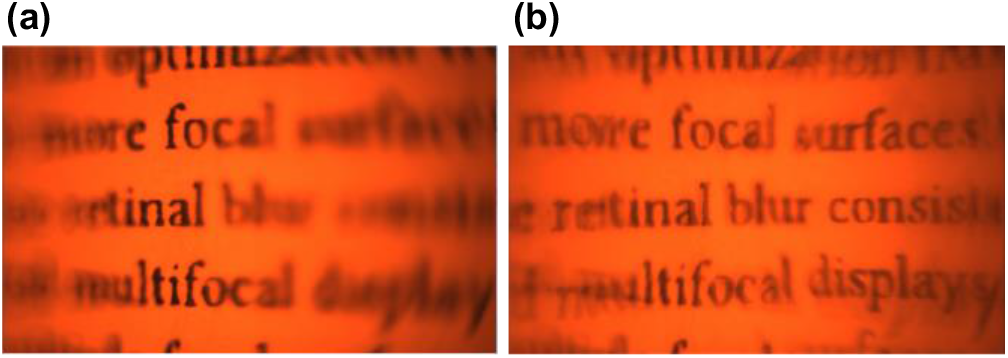
Reconstruction over a slanted planar surface. (a)-(b) Without and with applying the proposed contour imaging.

We then demonstrate the time-gating in improving the image resolution of TD-DOT. A 3D printed cylinder and cubic phantom (white ABS material, diameter/length of 20 mm) are made with three voids of square, circular and elliptical shape. The scattering properties inside the voids are varied by filling them with solutions of different concentrations of water and milk. To make the phantom contours at all rotation angles to fall within the depth of field, which is evaluated to be 8 mm under a circle of confusion of 1 mm, both phantoms’ center are carefully aligned with the rotation center. This allows good imaging quality without applying the proposed contour imaging. Figure 4 shows, from left to right, the phantom photographs and the reconstructed images with and without applying a 300 ps time gate on the full temporal waveform. For both phantoms, the reconstructed square and circular voids, which have smaller scattering coefficients than the background, are significantly larger than their ground truths. In contrast, the time-gated reconstruction results show better morphological agreement with the ground-truths and renders a clearer boundary. The estimated imaging resolution using 300 ps is about 2 mm and could be further improved by using a shorter time-gate. However, given the effective laser power of 20 μW, shorter time-gate yields noisier temporal waveform that ultimately compromise reconstruction stability and quality.

**Fig. 4.**
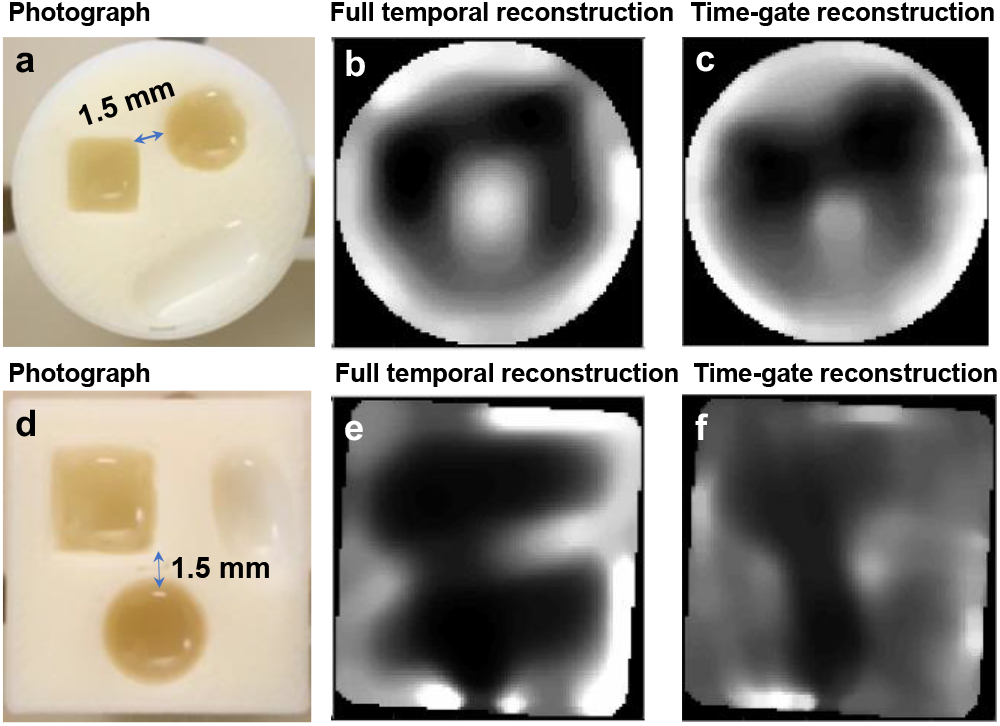
TD-DOT imaging results for a cylindrical and a cubic phantom. From left to right: photograph, reconstruction using the full temporal waveform and time-gated waveform with a duration of 300 ps. (a)-(c): cylindrical phantom. (d)-(f): cubic phantom.

The importance of maintaining the phantom in focus at all angular positions by the proposed ultrafast contour imaging is demonstrated in Fig. 5(a)-(f), where both the square phantoms are misaligned with the rotation center, leading to a maximum defocus error ±5 mm (outside the depth of field) at some rotation angles. As in previous experiments, the time-gate is 300 ps and the voids are filled with milk solution of different concentrations. Compared with the reconstructed images by the proposed ultrafast contour imaging method in (c) and (f), the defocus error in data acquisition causes the reconstructions in (b) and (e) to show significantly more artefacts and a less well-defined boundary of the voids.

**Fig. 5.**
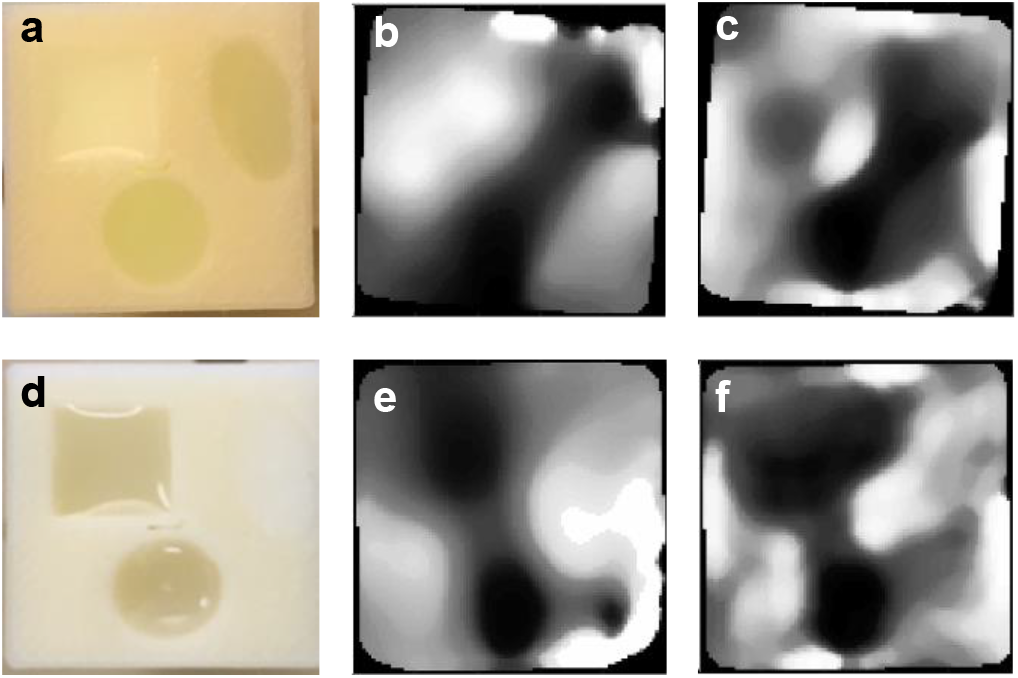
TD-DOT reconstruction of the square phantoms with and without applying the contour imaging method. From left to right: photograph, reconstruction without (b and e) and with (c and f) applying the contour imaging method, using time-gated waveforms with a duration of 300 ps.

Lastly, we demonstrate the proposed ultrafast contour imaging method for small animal DOT imaging under a protocol approved by the IACUC committee of University of Illinois at Urbana-Champlain. A six week old nude mouse was anesthetized with 100 mg/kg ketamine and 10 mg/kg xylazine administered IP using a sterile a 25-gauge needle and then mounted on a 3D printed holder for TD-DOT imaging. Fixed at an upright position at its native geometry, the size of the chest part is about 30 mm, which is slightly larger than those in previous studies. The photograph of the chest part at one angle was captured by the depth camera and is shown in Fig. 6(a). The corresponding time-resolved DOT signals across the slit of the streak camera before pixel binning is given in (b), with a time window of 5 ns. Despite of the low effective laser power and a larger trunk size, the large detection NA allows efficient light collection over the mouse native geometry and enables the reconstructed image in (c) to clearly identify the lung shapes in the chest region. Notably, the obtained chest DOT image agrees well with that obtained by x-ray CT in the literature.

**Fig. 6.**
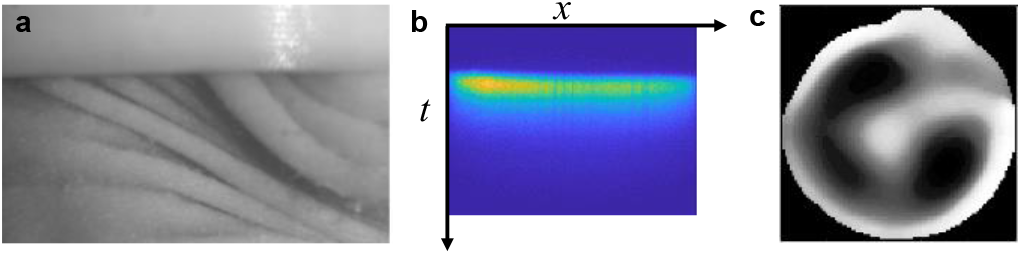
TD-DOT imaging of the chest part of a mouse. (a) Photograph of the mouse chest part at one rotation angle. (b) One representative spatiotemporal *x-t* DOT data. (c) Reconstructed image of the mouse chest part. The size of the mouse contour is about 30 mm.

While the current system with a detection NA of 0.25 still provides a depth of field of 8 mm at object side and is generally enough for small animal imaging, the depth of field will be reduced to about 2.5 mm once scaling the NA to 0.5, making it impractical for non-contact DOT imaging even when the object’s center is aligned with the mechanical rotation center. In our current system, scaling to a larger detection NA is challenging without compromising the field of view: the implementation using a beam splitter and a reflective SLM restricts very short back focal length to be used for the imaging lens. However, this restriction could be easily lifted by using a transmission SLM at the cost of a lower pixel resolution. There are also a few other techniques that can extend the depth of field while maintaining a large imaging NA. In general, all these techniques employs wavefront coding (by a SLM) to modulate the imaging point spread function (PSF), which can be depth-invariant like that produced by a cubic phase plate [25] or depth-variant such as the double-helix PSF [26], such that the image could be effectively deconvolved over an extended 3D imaging volume. The need of image deconvolution mandates the access to a properly sampled 2D image, thus precluding 1D ultrafast detectors like streak camera to be used. The proposed contour imaging method, in contrast, can imaging over a curved surface (effectively extend the depth of field) without the need of post-processing, making it more appealing for ultrafast detectors that usually come with a low pixel resolution.

In conclusion, we proposed a novel ultrafast contour imaging method, which can image over a curved surface without defocus error and thus enables imaging non-planar objects in their native geometry while maintaining a large detection NA for efficient light collection. When coupled with parallel ultrafast detectors such as streak cameras or intensified CCD, it facilitates time-domain DOT imaging with a high density data set and a shorter time-gating for improving the image resolution without compromising reconstruction stability. Further improvements on the system with a larger effective laser power and detection NA are potential to achieve sub-millimeter resolution for small animal imaging by applying a time-gate shorter than 200 ps.

## Funding

National Science Foundation (NSF) (CAREER 1652150); National Institute of Health (R01EY029397, R35GM128761).

## Acknowledgment

We thank the Biophotonic Imaging Group at University of Illinois at Urbana-Champaign for making the femtosecond laser available for this research study.

## Disclosures

The authors declare no conflicts of interest.

## Notes

### Competing Interest Statement

The authors have declared no competing interest.

## References

1. Y. H. M.d and Y. Yamada, JBO 21, 091312 (2016).

2. A. P. Gibson, J. C. Hebden, and S. R. Arridge, Phys. Med. Biol. 50, R1 (2005).

3. S. R. Arridge, Inverse Problems 15, R41 (1999).

4. S. D. Konecky, G. Y. Panasyuk, K. Lee, V. Markel, A. G. Yodh, and J. C. Schotland, Opt. Express 16, 5048 (2008).

5. H. Jiang, (CRC Press, 2018).

6. F. Leblond, H. Dehghani, D. Kepshire, and B. W. Pogue, J. Opt. Soc. Am. A, JOSAA 26, 1444 (2009).

7. K. Chen, L. T. Perelman, Q. Zhang, R. R. Dasari, and M. S. Feld, JBO, JBOPFO 5, 144 (2000).

8. M. J. Niedre, R. H. de Kleine, E. Aikawa, D. G. Kirsch, R. Weissleder, and V. Ntziachristos, Proceedings of the National Academy of Sciences 105, 19126 (2008).

9. G. M. Turner, A. Soubret, and V. Ntziachristos, Medical Physics 34, 1405 (n.d.).

10. C. Liu, A. K. Maity, A. W. Dubrawski, A. Sabharwal, and S. G. Narasimhan, in 2020 IEEE International Conference on Computational Photography (ICCP) (2020), pp. 1–12.

11. E. E. Graves, J. Ripoll, R. Weissleder, and V. Ntziachristos, Medical Physics 30, 901 (n.d.).

12. M. D. Wheelock, J. P. Culver, and A. T. Eggebrecht, Review of Scientific Instruments 90, 051101 (2019).

13. B. R. White and J. P. Culver, JBO 15, 026006 (2010).

14. A. T. Eggebrecht, S. L. Ferradal, A. Robichaux-Viehoever, M. S. Hassanpour, H. Dehghani, A. Z. Snyder, T. Hershey, and J. P. Culver, Nature Photonics 8, 448 (2014).

15. B. W. Zeff, B. R. White, H. Dehghani, B. L. Schlaggar, and J. P. Culver, PNAS 104, 12169 (2007).

16. S. M. Liao, S. L. Ferradal, B. R. White, N. M. Gregg, T. E. Inder, and J. P. Culver, JBO 17, 081414 (2012).

17. R. B. Schulz, J. Peter, W. Semmler, C. D’Andrea, G. Valentini, and R. Cubeddu, Opt. Lett., OL 31, 769 (2006).

18. N. Deliolanis, T. Lasser, D. Hyde, A. Soubret, J. Ripoll, and V. Ntziachristos, Optics Letters 32, 382 (2007).

19. E. Lapointe, J. Pichette, and Y. Bérubé-Lauzière, Review of Scientific Instruments 83, 063703 (2012).

20. N. Matsuda, A. Fix, and D. Lanman, ACM Trans. Graph. 36, 86:1 (2017).

21. E. M. C. Hillman, J. C. Hebden, F. E. W. Schmidt, S. R. Arridge, M. Schweiger, H. Dehghani, and D. T. Delpy, Review of Scientific Instruments 71, 3415 (2000).

22. M. Schweiger and S. R. Arridge, Appl. Opt., AO 37, 7419 (1998).

23. S. R. Arridge, M. Schweiger, M. Hiraoka, and D. T. Delpy, Med. Phys. 20, 299 (1993).

24. M. Schweiger, S. R. Arridge, and I. Nissilä, Phys. Med. Biol. 50, 2365 (2005).

25. E. R. Dowski and W. T. Cathey, Applied Optics 34, 1859 (1995).

26. Z. Wang, Y. Cai, Y. Liang, X. Zhou, S. Yan, D. Dan, P. R. Bianco, M. Lei, and B. Yao, Biomed. Opt. Express, BOE 8, 5493 (2017).

